# Seizure epicenter depth and translaminar field potential synchrony underlie complex variations in tissue oxygenation during ictal initiation

**DOI:** 10.1101/172148

**Authors:** Samuel S. Harris, Luke W. Boorman, Aneurin J. Kennerley, Paul S. Sharp, Chris Martin, Peter Redgrave, Theodore H. Schwartz, Jason Berwick

## Abstract

Whether functional hyperemia during epileptic activity is adequate to meet the heightened metabolic demand of such events is controversial. Whereas some studies have demonstrated hyperoxia during ictal onsets, other work has reported transient hypoxic episodes that are spatially dependent on local surface microvasculature. Crucially, how laminar differences in ictal evolution can affect subsequent cerebrovascular responses has not been thus far investigated, and is likely significant in view of possible laminar-dependent neurovascular mechanisms and angioarchitecture. We addressed this open question using a novel multi-modal methodology enabling concurrent measurement of cortical tissue oxygenation, blood flow and hemoglobin concentration, alongside laminar recordings of neural activity, in a urethane anesthetized rat model of recurrent seizures induced by 4-aminopyridine. We reveal there to be a close relationship between seizure epicenter depth, translaminar LFP synchrony and tissue oxygenation during the early stages of recurrent seizures, whereby deep layer seizures are associated with decreased cross laminar synchrony and prolonged periods of hypoxia, and middle layer seizures are accompanied by increased cross-laminar synchrony and hyperoxia. Through comparison with functional activation by somatosensory stimulation and graded hypercapnia, we show that these seizure-related cerebrovascular responses occur in the presence of conserved neural-hemodynamic and blood flow-volume coupling. Our data provide new insights into the laminar dependency of seizure-related neurovascular responses, which may reconcile inconsistent observations of seizure-related hypoxia in the literature, and highlight a potential layer-dependent vulnerability that may contribute to the harmful effects of clinical recurrent seizures. The relevance of our findings to perfusion-related functional neuroimaging techniques in epilepsy are also discussed.

## 1. Introduction

Understanding the physiological cascade of blood flow and metabolism during epileptic seizures can provide insights into the pathophysiology and pathological consequences of such events, and enhance the interpretation of diagnostic perfusion-related neuroimaging techniques in epilepsy. There is ongoing debate about whether the supranormal metabolic demands made by epileptic seizures are adequately met by local delivery of oxygenated blood through functional hyperemia. The notion that cerebral perfusion changes might not sustain these intense energy demands and lead to ischemia-hypoxia and neurodegeneration was advocated by a number of early studies (Spielmeyer, 1930; Davis et al., 1944; Cooper et al., 1966; Plum et al., 1968; Dymond and Crandall, 1976). However, this hypothesis fell largely out of favor with subsequent reports of hyperoxia during repetitive seizures with blood flow exceeding metabolic demand (Franck et al., 1985; Ingvar, 1986), and excitotoxic-related neuronal injury in the absence of hypoxia (for review see Meldrum, 2002). Notwithstanding, recent work has begun to come full-circle and provided renewed support for cerebrovascular dysfunction as a parallel phenomenon in epilepsy. Transient decreases in brain tissue oxygenation have recently been observed in the seizure onset zone during onset of ictal-like discharges (Bahar et al., 2006; Zhao et al., 2009), that are intensified in brain regions distal to arteries and arterioles, and can even serve to predict seizure duration (Zhang et al., 2015; Zhang et al., 2017). Other work has also provided support for ischemia-hypoxia during ictal events by demonstrating that ictal neurodegeneration is spatially coupled to abnormal capillary blood flow (Leal-Campanario et al., 2017), in keeping with computational models suggesting that altered capillary flow patterns can adversely affect oxygen extraction efficacy (Jespersen and Østergaard, 2012). These findings have therefore provided important insights into neurovascular ictal dynamics and the spatial role of microvasculature on these responses in the superficial plane of the cortex, although the question why tissue oxygenation deficits during ictal events have not been routinely observed remains unanswered. One relevant, but neglected, line of inquiry concerns the influence of the differential laminar evolution of ictal activity on accompanying hemodynamic and tissue oxygenation dynamics, given depth-specific variations in angioarchitecture (Masamoto et al., 2004; Blinder et al., 2013), tissue oxygen tension (Feng et al., 1988), and neurovascular coupling (Poplawsky et al., 2015). Laminar differences in seizure initiation and propagation could therefore significantly modulate ensuing cortical tissue oxygenation, thereby providing a mechanistic basis for variability in seizure outcomes and seizure-related oxemic observations in the literature. Unfortunately, previous methodological approaches have been unsuitable and unable to test this prospect directly.

Elucidating the extent to which the relationship between hemodynamic changes and underlying neural activity, known as neurovascular coupling, is conserved during epileptic seizures, compared to normal conditions, is also essential to the correct application of diagnostic perfusion-related neuroimaging techniques, such as blood oxygen level dependent (BOLD) functional magnetic resonance imaging (fMRI). BOLD-fMRI signals are typically interpreted in terms of underlying neural activation within the framework of a general linear model (Logothetis et al., 2001) that is invariant across health and disease, and are being increasingly employed in combination with electroencephalogram (EEG) recordings to localize hemodynamic correlates of electrographic ictal activity (Salek-Haddadi et al., 2002; Tyvaert et al., 2008). Furthermore, ascertaining the stability of the relationship between blood flow and volume is central to refining assumptive estimates of cerebral metabolic rate of oxygen (CMRO_2_) transients from human ‘calibrated’ fMRI (Davis et al., 1998; Kida et al., 2007), which can provide more faithful measures of neural activity compared to BOLD signals, and are therefore of potential value to ictal-fMRI studies. Nevertheless, whether normal neurovascular and blood-volume coupling is conserved during epileptic seizures remains to be fully clarified.

Accordingly, we have sought to address these unresolved questions through development of a novel multi-modal methodology that enables concurrent measurement of tissue oxygenation, blood flow and total hemoglobin concentration, alongside laminar neural activity, in the urethane anesthetized rat. We aimed to examine the role of laminar-dependent initiation and propagation of recurrent acute neocortical seizures, induced by 4-aminpyridine, on ensuing cerebrovascular responses. Further, through comparison to functional activation by whisker stimulation and hypercapnia, we endeavored to describe the response dynamics of all variables along a continuum of brain activation, and interrogate the stability of neural-hemodynamic and blood flow-volume coupling during ictal-like discharges.

## 2. Methods

### 2.1 Animal preparation and surgery

All procedures were conducted with approval from the UK Home Office under the Animals (Scientific Procedures) Act of 1986. Female hooded Lister rats (total N=12 weighing 250–350g) were kept in a 12-hr dark/light cycle environment at a temperature of 22°C, with food and water *ad libitum.* Animals were anesthetized with intraperitoneal injection of urethane (1.25 g/kg) and anesthetic depth was monitored throughout surgery and experiments. Urethane anesthesia preserves excitatory/inhibitory synaptic transmission (Sceniak and MacIver, 2006) and neurovascular coupling (Berwick et al., 2008), as well as providing persistent anesthesia reminiscent of natural sleep (Pagliardini et al., 2013). Core/body temperature of animals was maintained at 37°C through use of a homoeothermic blanket and rectal probe (Harvard Apparatus). Room temperature was thermostatically controlled at 19.1^o^C. Animals were tracheotomized and artificially ventilated with medical air. Blood-gas and end-tidal CO_2_ measurements were taken to adjust ventilator parameters and maintain the animal within normal physiological limits during experiments. The left femoral artery and vein were cannulated to allow the measurement of arterial blood pressure and phenylephrine infusion (0.13-0.26 mg/h to maintain normotension between 100 and 110 mmHg), respectively (Berwick et al., 2005). The animal was secured in a stereotaxic frame throughout experimentation and the skull overlying the right parietal cortex thinned to translucency so as to expose the somatosensory cortex. A layer of cyanoacrylate glue was subsequently applied over this region to reduce optical specularities from the brain surface during imaging and preserve skull integrity.

### 2.2 Two-Dimensional Optical Imaging Spectroscopy

Two-dimensional optical imaging spectroscopy (2D-OIS) was used to produce 2D images over time of total hemoglobin (Hbt) concentration, using a heterogeneous tissue model as described previously (Berwick et al., 2005; Berwick et al., 2008; Kennerley et al., 2009). Briefly, the cortex was illuminated at four different wavelengths (495±31 nm, 559±16 nm, 575±14 nm and 587±9 nm FWHM) using a Lambda DG-4 high speed filter changer (Sutter Instrument Company, Novata, CA, USA) and image data recorded at 8Hz using a Dalsa 1M30P camera (Billerica, MA, USA, each pixel representing ~75μm^2^) synchronized to the filter switching (2Hz/wavelength). Data were subsequently subjected to spectral analysis consisting of a path length scaling algorithm (PLSA) employing a modified Beer–Lambert law in conjunction with a path-length correction factor for each wavelength used, based on Monte Carlo simulations of light transport through tissue (Wang et al., 1995; Berwick et al., 2005; Berwick et al., 2008). Baseline concentration of Hbt was estimated at 104μmol/L (Kennerley et al., 2009). The somatosensory barrel cortex was localized prior to experiments to guide implantation of a multi-channel electrode and multi-sensor probe. Here, the left (contralateral) whisker pad was briefly stimulated electrically using subcutaneous electrodes (30 trials, 2s, 5Hz, 1.2mA, 0.3ms pulse width) and resultant 2D-OIS data averaged and subjected to the above spectral analysis. Spatiotemporal changes in Hbt were analyzed using statistical parametric mapping (SPM) which produced a Z-score activation map that was used to localize the cortical region activated by whisker stimulation (as described previously, Harris et al., 2013).

### 2.3 Multi-sensor probe recordings

A multi-modal sensor comprising of laser Doppler flowmetry (LDF) and luminescence-based oxygen probe (450μm diameter, Oxford Optronix, UK) was used to obtain concurrent measures of cerebral blood flow (CBF) and tissue oxygenation (tpO_2_), respectively (Trübel et al., 2006). The laser Doppler probe provided a measure of relative changes in CBF, while the luminescence probe provided an absolute measure of tpO_2_ (±0.1mmHg, 0.5mm diameter sphere sampling volume), respectively. A 750nm low-pass filter was attached to the 2D-OIS camera lens to prevent cross-talk from the LDF probe operating at 830±10nm, and the transmission curve of the filter accounted for during spectral analysis. The multi-sensor probe was attached to a stereotaxic holder and inserted into the somatosensory ‘whisker barrel’ cortex under microscope guidance to a depth of 500μm (layer 4) (Figure 1A). Signals were amplified and digitized using a 2-channel OxyFlo Pro system (Oxford Optronix, UK) and recorded using a CED Power 1401 and Spike2 software (Cambridge Electronic Design, Cambridge, UK). Sensor data were recorded continuously during each experiment and sampled at 1Hz.

**Figure 1:**
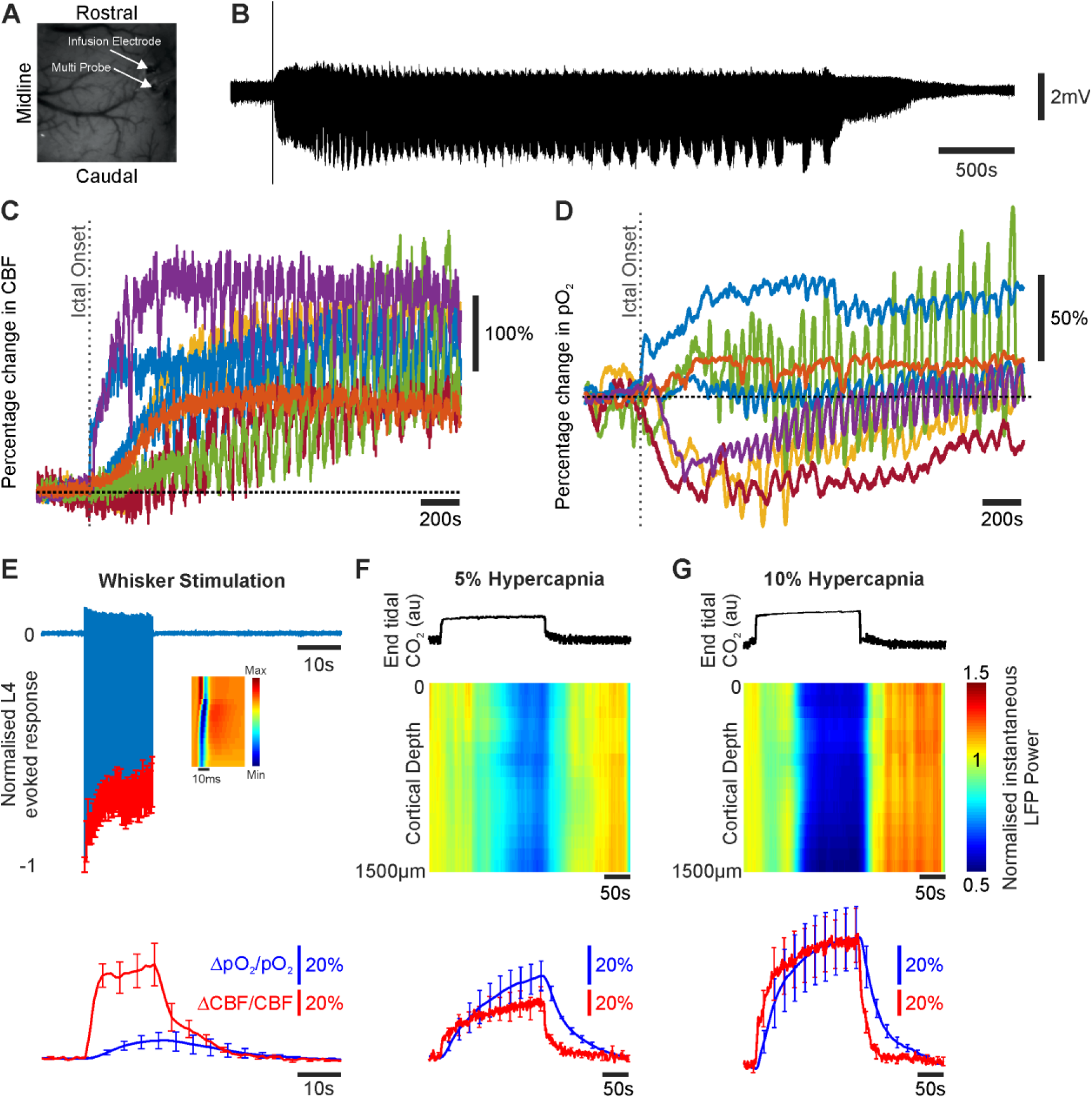
Neural, CBF and oxygenation changes during recurrent acute neocortical seizures, sensory stimulation and graded hypercapnia. A) Digital image of cortical surface indicating implantation site of multi-laminar electrode and multi-sensor for CBF and tissue oxygenation recordings in an example animal. B) Example LFP trace of recurrent seizure activity taken from 1500μm depth at 4AP infusion site. C) Percentage CBF changes in all animals (N=7) during first 2500s of recurrent seizure activity. D) Marked variability in percentage tpO_2_ changes in all animals (N=7) during first 2500s of recurrent seizure activity (color coded as in C). E) Normalized and averaged LFP layer 4 sink responses to 16s whisker stimulation (1.2mA, 5Hz, N=5, top) and accompanying averaged CBF (red) and tpO_2_ (blue) percentage changes during stimulation (bottom, N=5). Inset shows a representative laminar LFP response profile to first pulse in stimulation train (averaged across trials). F) Example end tidal CO_2_ trace (top), normalized instantaneous LFP power as a function of depth (middle, averaged across animals, N=5), and averaged CBF (red) and tpO_2_ (blue) percentage changes (bottom, N=5), during 5% hypercapnia. G) Same as in F but for 10% hypercapnia.

### 2.4 Laminar electrophysiology

A multi-channel microelectrode (16 channels with 100μm spacing, site area 177μm^2^, 1.5–2.7 MΩ impedance; Neuronexus Technologies, Ann Arbor, MI, USA) was placed in a stereotaxic holder and implanted into the right barrel cortex using a micro-manipulator and microscope to a depth of 1500μm (Harris et al., 2013; Boorman et al., 2015), and as close as possible to the multi-sensor probe (Figure 1A). During epilepsy experiments the multi-channel microelectrode was coupled to a fluidic probe (Neuronexus Technologies, Ann Arbor, MI, USA) to enable drug delivery. The microelectrode was connected to a preamplifier and data acquisition device (Medusa BioAmp/ RZ5, TDT, Alachua, FL, USA). Neural data were recorded continuously throughout each experiment and sampled at 6 kHz.

### 2.5 Experimental Paradigm

Multi-modal responses to whisker stimulation (N=5) were investigated using a 15 trial protocol, each delivering a 16-second train of electrical pulses (5 Hz, 1.2 mA intensity, 0.3ms pulse width) with an inter-trial interval of 70 seconds, to the right whisker pad following a 10-second baseline period (Harris et al., 2013). Hypercapnia experiments (N=5) were conducted after the stimulation protocol and consisted of seven interleaved trials of 190s duration with normocapnia on odd trials, and hypercapnia on trials 2, 4 and 6. No stimulation was delivered during hypercapnia experiments. Hypercapnia was induced in two sequential experimental runs where the fraction of inspired CO_2_ (FiCO_2_) was increased to either 5% (N=5) or 10% (N=5) (Jones et al., 2005). Hypercapnia trials for each condition were extracted with a 20s normocapnic baseline and 190s post-hypercapnic period. As hypercapnia has been demonstrated to have a potent anti-convulsant action (Tolner et al., 2011), we conducted acute seizure experiments in a separate group of animals (N=7). Fluidic microelectrodes were loaded with the potassium channel blocker 4-aminopyridine (4-AP, 15mM, 1μl, Sigma, UK) to induce recurrent focal seizure-like discharges in right barrel cortex (Harris et al., 2014a; Harris et al., 2014b). 4-AP was intra-cortically infused at a depth of 1500μm over a 5-minute period (0.2μl/min), using a 10μl Hamilton syringe and syringe pump (World Precision Instruments Inc., FL, USA) (Harris et al., 2014a; Harris et al., 2014b). Multi-sensor, electrophysiological and 2D-OIS measures were recorded concurrently for 7000s with 4-AP injection initiated following a 280s baseline period.

### 2.6 Data and statistical analysis

All data analysis was conducted using custom written Matlab^™^ scripts (Mathworks, Natick, MA, USA). Laminar neural recordings were bandpass filtered between 1 and 300Hz using a 300^th^ order finite impulse response filter and de-meaned to yield local field potential (LFP) data. Cross-laminar synchronization of neural activity was investigated through use of a mean phase coherence (MPC) algorithm, described in detail previously (Mormann et al., 2000; Schevon et al., 2007). The MPC algorithm was sequentially applied to pairs of LFP signals acquired from two different electrode sites along the multi-channel array (16 channels yielding 120 unique combinations of paired electrode data). Briefly, the instantaneous phase of each LFP signal was first calculated using the Hilbert transform, with instantaneous phase values of both signals projected onto the unit circle in the complex plane, and the mean phase coherence computed according to the expression (Mormann et al., 2000):

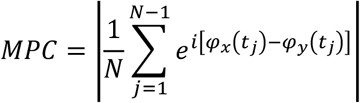

where *ϕ_x_* and *ϕ_y_* are the instantaneous phases of two input signals, *N* is the number of samples of the data set, and *t* is the time index. Euler’s formula was then used to transform the above equation, such that MPC is restricted to the interval [0,1], where MPC=0 denotes complete asynchrony and MPC=1 represents strict phase locking between both signals (Mormann et al., 2000). Importantly, MPC is not dependent on the amplitude of the input signals, and is insensitive to constant phase delays between two signals such as that arising due to neural transmission latency between two recording locations (Mormann et al., 2000; Schevon et al., 2007). Synchrony between paired LFP signals was obtained by computing the MPC in 200ms time-bins, applying a 5-point moving average filter, and downsampling the resultant timeseries to a sampling frequency of 1Hz (with use of an 8^th^ order lowpass Chebyshev Type I infinite impulse-response filter to minimize aliasing). Each MPC timeseries was normalized to its mean MPC value during the 280s baseline period prior to 4-AP infusion. LFP synchrony between different cortical layers was examined by grouping and averaging normalized MPC data relevant to the source-target cortical laminae of interest, defined according to cortical depth; supragranular (SG < 400μm), granular (GR, 400 - 700μm), infragranular (IG, 800 - 1100μm) and deep infragranular (dIG, 1200 - 1500μm).

Cerebral metabolic rate of oxygen (CMRO_2_) was calculated from CBF and tissue oxygen tension (tpO_2_) measurements according to the following relationship (Gjedde, 2005):

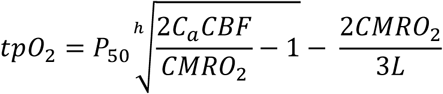

where tpO_2_ is the tissue oxygen tension and *P_50_* and *h* are the half-saturation tension and Hill coefficient of the oxygen–hemoglobin dissociation curve, 36mmHg and 2.7, respectively. *C_a_* is the arterial oxygen concentration (8μM/ml). The effective diffusion coefficient of oxygen in brain tissue, *L*, was calculated for each experiment using trial-averaged baseline tpO_2_ values, and baseline values of CBF and CMRO_2_ previously reported in rats by our laboratory and others, 47ml/100g/min (He et al., 2007) and 219μM/100g/min (Zhu et al., 2002), respectively. *L* was found to be 7.27±0.89μM/100g/min/mmHg (N=22 separate stimulation, hypercapnia and seizure experiments).

Time-series of Hbt responses during the different experimental conditions were extracted by selecting a region of interest (ROI) over acquired Hbt image data. As propagatory patterns during seizures can be highly variable and dynamic (Ma et al., 2012), we elected to manually position a ROI such that it was immediately adjacent to the multi-sensor probe and depth electrode, and captured a well-mixed vascular compartment (i.e. arterial, venous and parenchymal). The mean Hbt time-series of all pixels in the ROI were subsequently averaged for each animal and normalized to the baseline period prior to 4-AP infusion.

Since evoked cortical responses to stimulation exhibit a great deal of stability and reproducibility due to the fine control of stimulation parameters, Hbt image data acquired during each animal’s sensory stimulation condition were averaged over the period 0-20s after stimulus onset, and pixels possessing >80% of the maximum change in evoked Hbt automatically selected. The mean Hbt time-series of all pixels within the ROI were then extracted for whisker stimulation and hypercapnia conditions in turn. Hbt data from each animal during stimulation and graded hypercapnia were normalized to the mean baseline value across trials in each respective condition.

Recurrent seizure onset and offset in electrophysiological recordings was estimated by conducting power spectrum analysis on broadband field potential data from the deepest electrode channel (i.e. corresponding to the 4-AP infusion site at 1500μm, representative example shown in Figure 1A) using a Gabor transform. This consisted of short-term Fourier transform and 1s Gaussian window function applied sequentially to the entire timeseries (Harris et al., 2014b), and identifying the time-points where total broadband power first exceeded, and latterly fell below, 5 standard deviations above the mean baseline (i.e. 280s period prior to 4-AP infusion). Recurrent seizure duration was 5325.9±825.1s (N=7) and the seizure initiation period was defined as a 300s time-window following ictal onset. The locus of maximum activation during recurrent seizures (i.e. seizure epicenter) was found by calculating the total instantaneous power (integral of squared magnitude) of broadband laminar field potential data during the entire ictal period, and identifying the electrode channel exhibiting the greatest change in magnitude. Seizure epicenters were located on average at a depth of 1143±113μm (N=7), consistent with cortical layer 5b and our previous work (Harris et al., 2014a). Seizure-related LFP response profiles were obtained by identifying depth-negative peak responses during ictal activity whose magnitude was less than 5 standard deviations below the mean baseline activity. LFP profiles were then inverted and normalized to the largest peak response in each animal.

The layer 4 current sink in laminar neural recordings was localized by averaging the whisker evoked response to the first stimulus pulse across trials and identifying the electrode channel associated with the earliest robust negative field potential deflection; in this study the L4 current sink was located 580±111μm (N=5) below the cortical surface. Trial-by-trial Layer 4 LFP data were then extracted for each animal and normalized such that the peak (depth-negative) response evoked by the first pulse of the stimulation train was unity across trials (Harris et al., 2013) (representative animal shown in Figure 1E). Stimulation-related LFP response profiles were subsequently extracted by identifying the peak response to each electrical pulse in the stimulation train. Electrophysiological changes during hypercapnia were assessed by computing the instantaneous power of laminar LFP data and averaging over 1s time-bins to match the sampling frequency of the multi-sensor probe. Instantaneous power data from each electrode channel was normalized to the mean baseline value during the three trials in each hypercapnia condition.

In order to account for differing data ranges across experimental conditions, trial-by-trial relationships between variables during stimulation, hypercapnia and seizures were analyzed following standardization of the data using z-scores (i.e. with mean 0 and standard deviation 1). Timeseries are presented as a percentage change from baseline, and errors are given as the standard error from the mean (SEM), unless otherwise stated. Linear regression models were fitted using ordinary least squares, and differences between regression slopes examined using analysis of covariance with Tukey-Kramer correction for multiple comparisons. A 1-tailed Wilcoxon signed rank test was used to test significant changes in individual variables, and significant differences between conditions examined using a Kruskal-Wallis test with Tukey-Kramer correction for multiple comparisons. Exploratory correlation analyses involving micro-array data were corrected for using the Benjamini and Yekutieli (2001) procedure for controlling the false discovery rate (FDR) that is suitable for all forms of test-statistic dependency. Multiplicity-adjusted p values < 0.05 were defined as statically significant, with p < 0.01 deemed to be highly significant.

## 3. Results

### 3.1 Complex tissue oxygenation changes despite robust perfusion increases during early stages of recurrent seizures

The mean baseline oxygenation across animals prior to experimentation was 22.1±3.4mmHg (N=12), consistent with previous findings at cortical depths of 400-600μm in rats with intact skull preparations (Feng et al., 1988). Recurrent seizure activity was associated with considerable increases in CBF (maximal CBF: 258.8±35.7%, N=7, Figure 1C, 2A), that exceeded CBF increases during sensory stimulation (83.2±12.5%, N=5) and hypercapnia (50.2±10.3%, 113±23.4%, 5% and 10% FICO_2_ respectively, N=5 in both cases) (Figures 1E-F bottom panels, 2A). However, tPO_2_ changes were highly variable during the initial stages of recurrent seizure activity (Figure 1D), such that 3/7 animals exhibited a pronounced decrease immediately following seizure onset with subsequent recovery, with the remaining 4 animals exhibiting variable increases following seizure onset. Distinct polyphasic changes to individual ictal discharges, which recurred with short inter-seizure intervals, were also observed. In contrast, sensory stimulation and graded hypercapnia challenges elicited consistent increases in tPO_2_ (maximal tpO_2_: 10.4±4.7%, 42.8±7.6%, 71.4±16.2%, for stimulation, 5%, and 10% FICO_2_, respectively, N=5 in all cases, Figure 1E-G). As expected, estimates of mean CMRO_2_ changes, obtained from CBF and tpO_2_ recordings, provided confirmation of a dramatic and sustained increase in oxidative metabolism during recurrent seizures (mean CMRO_2_: 39.1±7.9%, N=7, see also Figure 2B) (e.g. Meldrum and Nilsson, 1976) that greatly outstripped those produced by normal functional activation during sensory stimulation (mean CMRO_2_: 8.9±2.9%, N=5, see also Figure 2B). Decreases in mean CMRO_2_ were also observed during graded hypercapnia, presumably as a result of decreases in LFP activity (Figure 1F-G, middle panels), although these were not found to be significant (p=0.91 and p=0.78, 5%, and 10% FICO_2_, respectively, N=5 in both cases, 1-sided Wilcoxon signed rank test, Figure 2B). Incidentally, this lends additional weight to the assumption that this popular calibration technique does not produce a significant change in CMRO_2_ (Davis et al., 1998; Kida et al., 2007).

**Figure 2:**
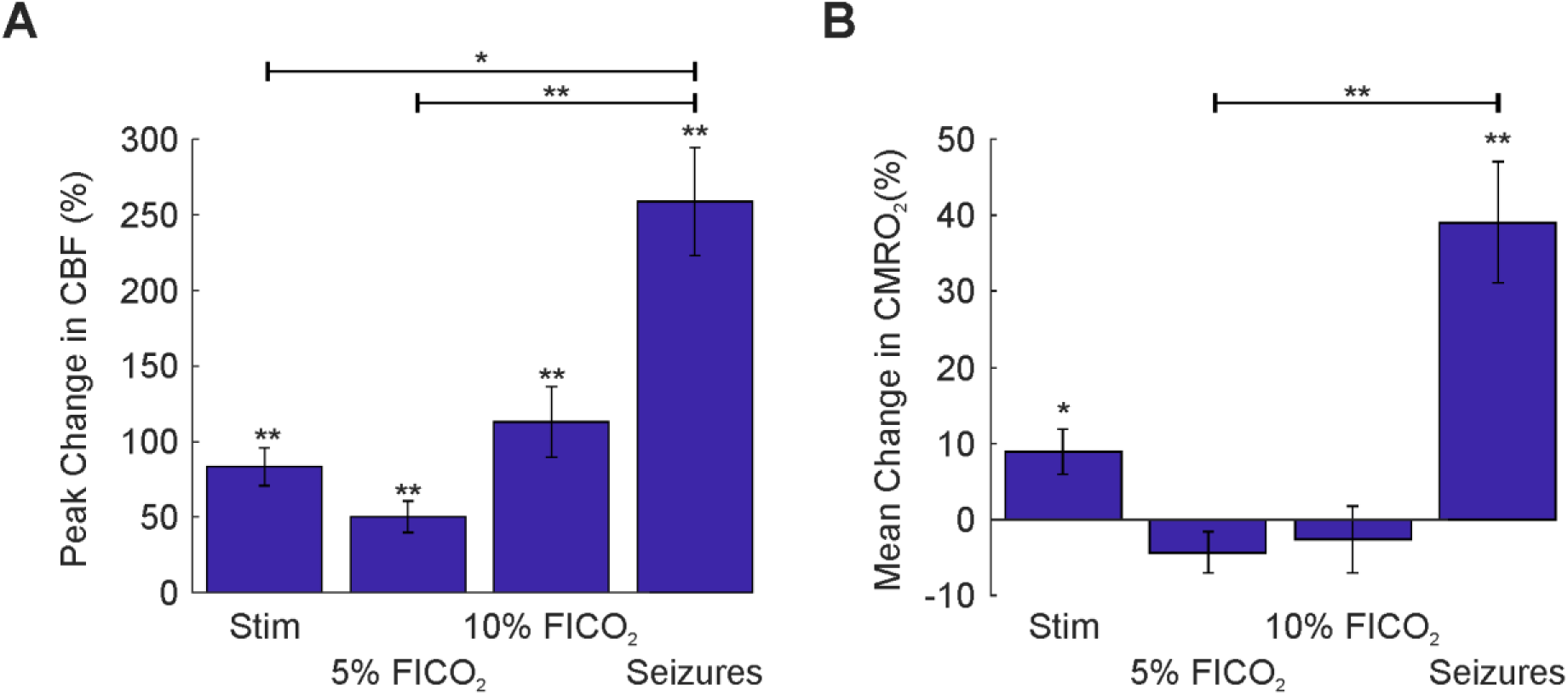
CBF and CMRO_2_ changes during sensory stimulation, graded hypercapnia and recurrent seizures. A) Peak percentage changes in CBF during whisker stimulation, 5% and 10% fraction of inspired CO_2_ (FICO_2_) and recurrent seizures. B) Mean percentage change in CBF during whisker stimulation (N=5), 5% and 10% fraction of inspired CO_2_ (FICO_2_, N=5 in each case) and recurrent seizures (N=7). Statistical comparisons made using Kruskal-Wallis tests with Tukey-Kramer correction. Individual comparisons made using 1-tailed Wilcoxon signed-ranks tests. ^*^P<0.05 and ^**^P<0.01. Error bars are SEM.

### 3.2 Variable tpO_2_ during recurrent seizures is related to laminar-dependent changes in LFP activation and synchrony

We next investigated candidate processes that might underpin the observed variability in tpO_2_ during recurrent seizures. CBF onset dynamics, which typically consisted of robust and rapid increases, was first ruled out as a possible driver of the observed tpO_2_ variability, since no significant correlation between both variables was observed during ictal onset (*p*=0.24, N=7) (example from two different animals with tpO_2_ changes of opposite polarity are shown in Figure 3A-B). Close inspection of laminar LFP waveforms, however, suggested that the vertical propagation of ictal activity differed between animals where tpO_2_ increased or decreased, with the latter associated with a comparatively decreased activation in upper cortical layers (example shown in Figure 3C-D). Mean phase coherence (MPC) analysis of LFP data from paired microelectrode channels was therefore conducted to assess seizure propagation though variations in laminar synchrony during seizure initiation. This revealed characteristic differences between cases where tpO_2_ increased or decreased during seizure onset (example shown in Figure 3E-F). Notably, the MPC between upper and lower cortical layers exhibited an increase during seizures accompanied by an increase in tpO_2_ (red areas in Figure 3E), whereas seizure onsets associated with a decrease in tpO_2_ showed the reverse (blue areas in Figure 3F).

**Figure 3:**
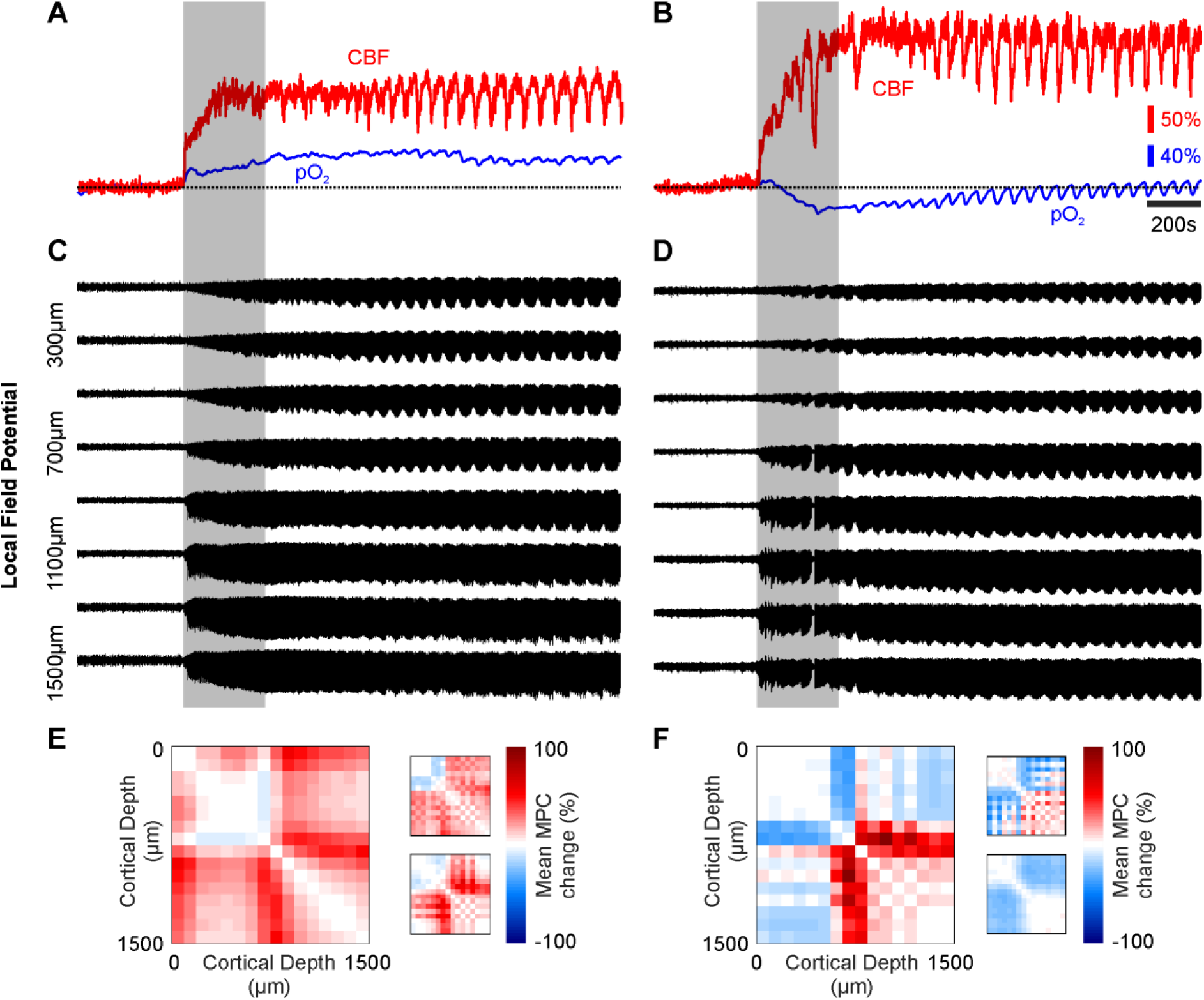
tpO_2_, CBF, laminar LFP activity and mean phase coherence (MPC) changes in two contrasting cases. A) Percentage changes in CBF (red) and tpO_2_ (blue) in an animal where tpO_2_ increases after seizure onset. B) Percentage changes in CBF (red) and tpO_2_ (blue) in a different animal where tpO_2_ decreases after seizure onset despite a marked rise in CBF. C) Associated laminar LFP recordings in same animal as A, showing strong activation in superficial depths. D) Laminar LFP recordings from same animal as B, showing comparatively weaker activation in superficial depths. E-F) Time-averaged normalized phase coherence matrices during seizure onset (300s, shaded bar in A-D) showing increases (red) and decreases (blue) in mean phase coherence (MPC) between electrode channel pairs. MPC matrices are symmetric about the diagonal. Left-hand matrix in E and F are from animals as in A, and B, respectively, with insets from two other representative animals with associated tpO_2_ increases and decreases, respectively.

The relationship between these coherence patterns and associated changes in tpO_2_ was confirmed using correlation analysis, which revealed MPC between supragranular and underlying laminae, and tpO_2_, to be significantly linearly correlated (SG-GR, Pearson’s r=0.91, p=0.02; SG-IG, Pearson’s r=0.91, p=0.02; SG-dIG, Pearson’s r=0.93, p=0.02; N=7, FDR corrected for multiple comparisons), at variance with other cortical laminae combinations (Figure 4A). Furthermore, cross-laminar changes in MPC during seizure onset, across all cortical depths studied, were also tightly coupled to changes in tpO_2_ (Pearson’s r=0.91, p=0.004, N=7, Figure 4B).

**Figure 4:**
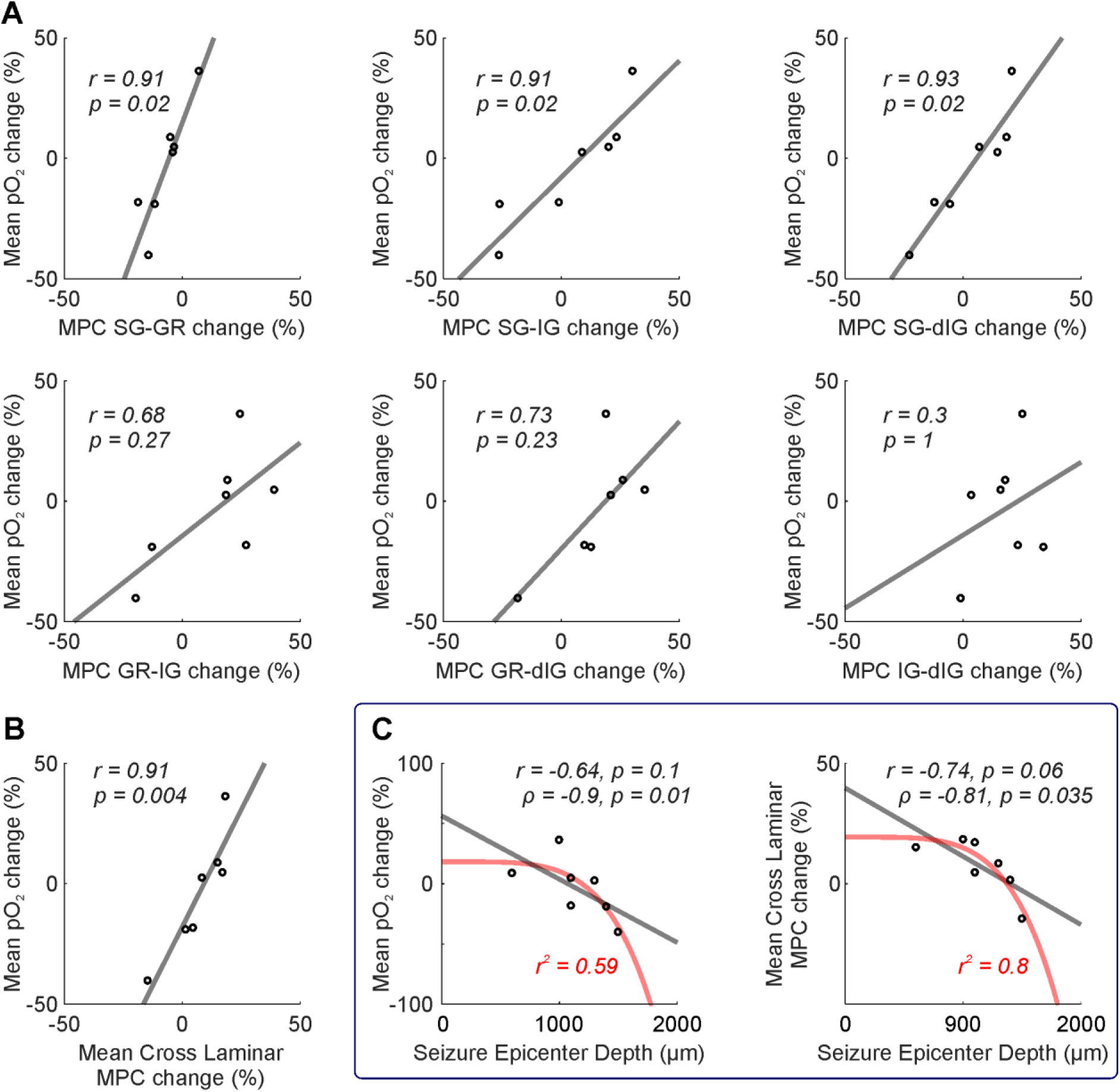
Inter-relationship between tpO_2_, laminar LFP synchrony and seizure epicenter depth. A) Laminar synchrony (MPC) spanning all laminar permutations (SG – supragranular, GR – granular, IG – infragranular, dIG – deep infragranular) against mean percentage change in tpO_2_ during seizure onset. Note significant linear correlations between MPC in supragranular and underlying laminae and tpO_2_ (top). B) Significant linear correlation between cross laminar synchrony (all laminae) and mean percentage change in tpO_2_ during seizure initiation. C) Significant inverse monotonic correlation between seizure epicenter depth and mean percentage change in tpO_2_ (left) and cross laminar synchrony (right) during seizure initiation. A-C) Linear (grey) and non-linear functions (red) fitted using least squares regression. Correlation coefficients and significance (FDR corrected for multiple comparisons), and r^2^ goodness-of-fit coefficient (red) for non-linear functions, given in each case.

Finally, we also found that the seizure epicenter depth was inversely monotonically related to mean tpO_2_ changes (Spearman’s rho=- 0.9, p=0.01, N=7, Figure 4C left panel), such that seizure epicenter depths >1200μm (consistent with L5b/6 border and beneath) were more likely to be associated with decreased tpO_2_ at seizure onset. Furthermore, seizure epicenter depth and cross-laminar changes in MPC during seizure onset were also negatively correlated (Spearman’s rho=- 0.9, p=0.01, N=7, Figure 4C right panel). Taken together, these results suggest that the epicenter depth of seizure activity and subsequent propagatory dynamics are closely interlinked, and play an important role in shaping tissue oxygenation changes during the early stages of recurrent seizure activity.

### 3.3 CBF-Hbt coupling is preserved during sensory stimulation, hypercapnia and recurrent seizures

Total hemoglobin concentration (Hbt), which is proportional to cerebral blood volume (CBV) under the assumption of a constant hematocrit, also increased markedly, albeit more slowly than CBF, during recurrent seizure activity (peak Hbt: 28.8±2.2%, N=7, Figure 5A top panel, 5B). Hbt responses to stimulation and graded hypercapnia also displayed archetypal features, with slower onset and return to baseline dynamics compared to CBF (maximal Hbt: 8.3±0.9%, 8.3±1.2%, 20.4±4%, for stimulation, 5%, and 10% FICO_2_, respectively, N=5 in all cases, Figure 5A bottom panels, 5B). Of note, no significant difference in peak Hbt was observed between 10% FICO_2_ and recurrent seizure conditions (p=0.38, Kruskal-Wallis test with post hoc Tukey Kramer analysis, Figure 5B), mirroring that seen with CBF (p=0.22, Kruskal-Wallis test with post hoc Tukey Kramer analysis, Figure 2A), and suggesting that higher levels of hypercapnia can induce perfusion-related responses comparable to recurrent seizures.

**Figure 5:**
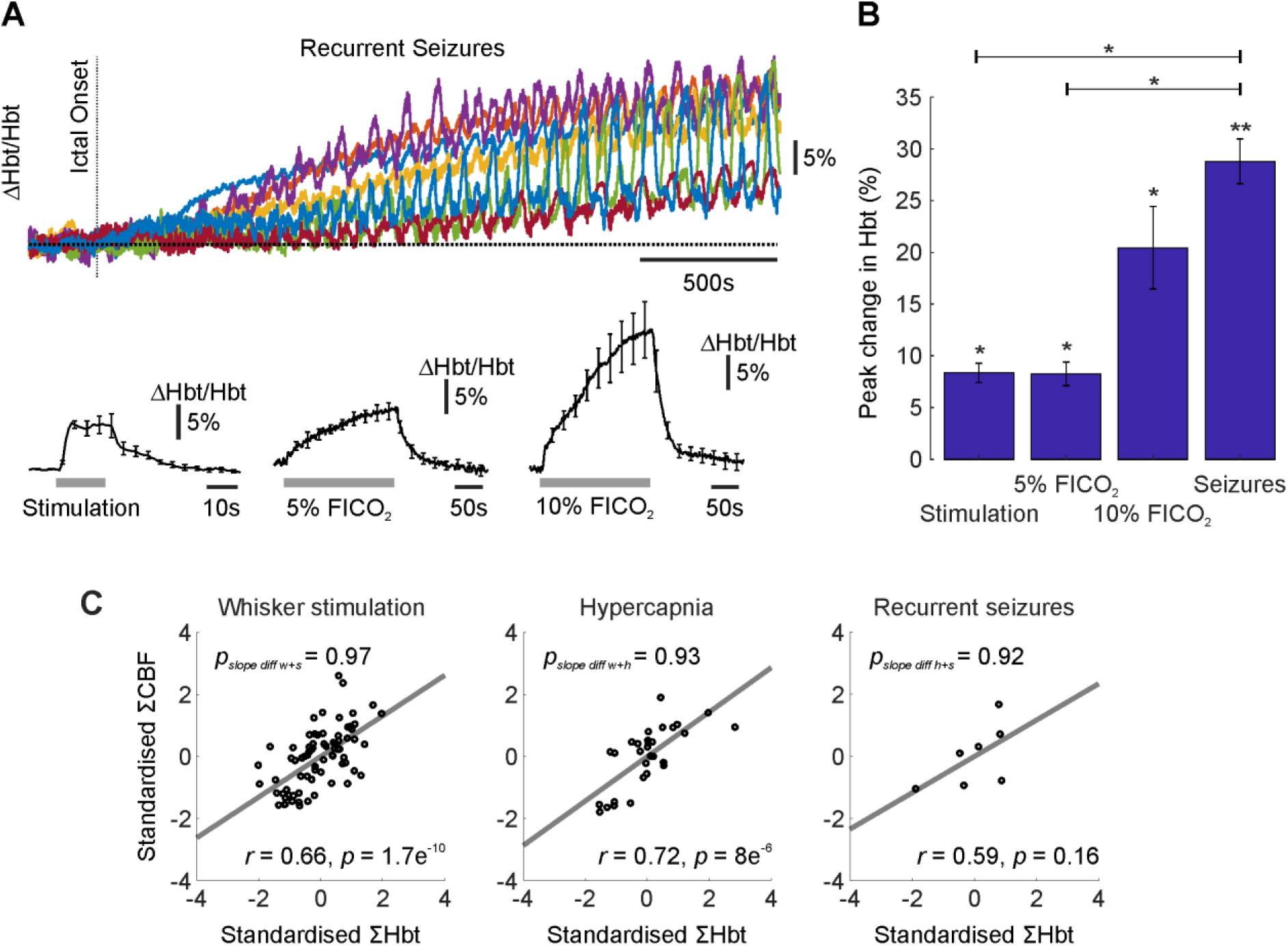
Hbt changes during recurrent seizures, sensory stimulation and graded hypercapnia, and Hbt-CBF coupling. A) Percentage changes in Hbt during first 2500s of seizure activity in all animals (top) and average during sensory stimulation and graded hypercapnia (bottom, N=5). B) Mean percentage change in Hbt during whisker stimulation (N=5), 5% and 10% fraction of inspired CO_2_ (FICO_2_, N=5 in each case) and recurrent seizures (N=7). A-B) Statistical comparisons made using Kruskal-Wallis tests with Tukey-Kramer correction. Individual comparisons made using 1-tailed Wilcoxon signed-ranks tests. ^*^P<0.05 and ^**^P<0.01. Error bars are SEM. C) Trial-by-trial analysis showing strong linear correlation between standardized Hbt and CBF during stimulation (‘w’, N=75), hypercapnia (‘h’, pooled across graded FICO_2_, N=30) and recurrent seizures (‘s’, N=7). Grey linear functions fitted using least squares regression. Correlation coefficients and significance given as insets in each case. Significant differences between all combinations of regression slopes were tested using analysis of covariance with T ukey-Kramer correction for multiple comparisons and given as insets in each case (top).

Trial-by-trial analysis was subsequently performed to assess the relationship between CBF and Hbt within and across experimental conditions. CBF-Hbt linear correlations were observed across experimental conditions (Figure 5C) that were highly significant in the case of stimulation and pooled hypercapnia (Pearson’s r=0.66, p < 0.01, N=75, and r=0.72, p <0.01, N=30, respectively), and approached significance with respect to recurrent seizures (Pearson’s r=0.59, p=0.16, N=7). Importantly, linear regression slopes were not significantly different between conditions (slope difference between whisker stimulation and seizures, p=0.99; slope difference between whisker stimulation and hypercapnia, p=0.93; and slope difference between hypercapnia and seizures, p=0.97), suggesting that a similar relationship between CBF-Hbt is preserved across a broad continuum of brain activation.

### 3.4 Neural-hemodynamic coupling is preserved during sensory stimulation and recurrent seizures

We then asked whether the relationship between neural activity and perfusion related measures was maintained across normal (whisker stimulation) and aberrant (recurrent seizures) functional activation. Trial-by-trial analysis revealed a significant linear correlation between summated L4 LFP activity and CBF during whisker stimulation (Pearson’s r=0.7, p < 0.01, N=75, Figure 6A, blue), and summated LFP responses in the ictal epicenter (site of maximal ictal activation) and CBF during recurrent seizures (Pearson’s r=0.81, p=0.01, N=7, Figure 6A, red). Linear regression slopes did not differ significantly between whisker stimulation and recurrent seizure conditions (p=0.63), although the latter did exhibit a slightly steeper gradient (β=0.7 and 0.86, respectively). Correspondingly, we also observed a significant linear correlation between L4 LFP activity and Hbt during whisker stimulation (Pearson’s r=0.39, p < 0.01, N=75, Figure 6B, blue), and LFP responses at the site of maximal ictal activation and Hbt during recurrent seizures (Pearson’s r=0.96, p < 0.01, N=7, Figure 6B, red). Yet again, no significant difference was found between regression slopes during whisker stimulation and recurrent seizures (p=0.22), with the latter displaying once more a steeper gradient relative to the former (β=0.39 and 0.9, respectively). Taken together, these results suggest that neural-hemodynamic coupling remains remarkably robust across normal and ictal forms of functional activation.

**Figure 6:**
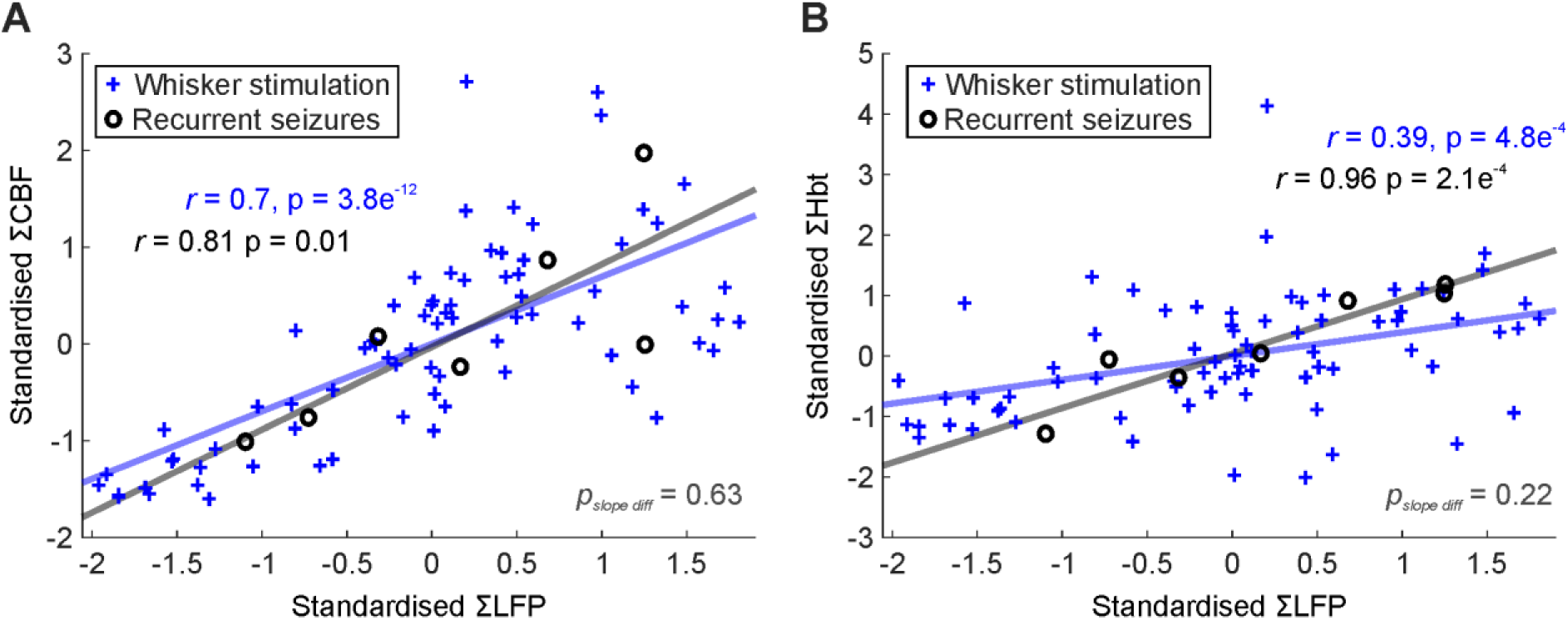
Conserved neural-hemodynamic coupling during functional activation by sensory stimulation and recurrent seizures. A) Trial-by-trial analysis of relationship between standardized LFP and CBF during sensory stimulation (blue crosses, N=75) and recurrent seizure activity (black open circles, N=7). B) Trial-by-trial analysis of relationship between standardized LFP and Hbt during sensory stimulation (blue crosses, N=75) and recurrent seizure activity (black open circles, N=7). A-B) Linear functions fitted using least squares regression with Pearson correlation coefficients and significance given as insets in each case. Significant differences between pairs of regression slopes were tested using analysis of covariance and given as insets in each case (bottom).

## 4. Discussion

The principal finding of this study is that of a previously unreported link between laminar LFP synchrony and depth-related ictal hypoxia. Notably, repetitive seizures with deep cortical epicenters were associated with prolonged periods of tissue hypoxia and reduced cross-laminar LFP synchrony, despite considerable increases in perfusion and preserved neural-hemodynamic, and CBF-Hbt, coupling. Our data suggest that functional hyperemic mechanisms at the pial-arteriole network level (to which our CBF and Hbt measures are biased) continue to operate normatively during repetitive seizures, but are insufficient to maintain normoxia during the onset of deep lying focal seizures, most likely as a result of local depth-dependent microvascular susceptibilities. These findings provide new insights into seizure-related cerebrovascular dysfunction and may be relevant to the interpretation of perfusion-related neuroimaging signals in epilepsy.

### 4.1 Relationship between cortical tissue oxygenation, depth of the ictal epicenter, and laminar LFP synchrony, during the onset of recurrent seizures

Layer 6 is believed to play a key role in cortical gain control and thus sensory processing, with optogenetic activation of L6 corticothalamic neurons in visual cortex found to suppress activity across all laminae through recruitment of translaminar fast-spiking inhibitory neurons (Olsen et al., 2012; Bortone et al., 2014; Kim et al., 2014). Recurrent seizures whose epicenter reside in this cortical layer may therefore be predisposed to being more spatially restrained by an intense overlying inhibitory surround (Prince and Wilder, 1967; Schevon et al., 2012), leading to an overall decrease in cross laminar neural synchrony and ‘functional disconnection’ from surrounding cortex (Warren et al., 2010). Functional isolation of seizure-generating cortex may prevent pathologically firing neurons in the ictal focus from being mollified by the surrounding network and allow these to develop hyper-intense excitatory states (Uhlhaas and Singer, 2006). It may also underpin the accompanying robust decrease in tpO_2_ during the initiation of these seizures. We have previously reported the presence of smaller, more transient, ‘dips’ in tissue oxygenation during onset of singular 4AP discharges in the ictal focus (Bahar et al., 2006; Zhao et al., 2009), mirroring that seen during human seizures (Cooper et al., 1966; Dymond and Crandall, 1976). Importantly, it has since been demonstrated that the magnitude of this dip, in the same seizure model, is positively correlated to the horizontal distance between the seizure focus and surface arteries and arterioles (Zhang et al., 2015; Zhang et al., 2017). This is presumably due to steep tpO_2_ gradients decreasing as a function of distance from such vessels, and rendering distal tissue comparatively more susceptible to hypoxia during metabolically demanding events (Sakadžić et al., 2010; Kasischke et al., 2011).

It is thus possible that recurrent seizures in deep laminae may also be vulnerable to similar heterogeneity in tissue oxygenation landscape in the depth dimension, given that the absence of a significant correlation between tissue oxygenation and CBF changes during seizure onset, as well as a preserved neural-hemodynamic relationship, argues against a perfusion-related cause for the observed fall in tpO_2_. Indeed, previous work in rat using open and closed skull preparations has shown that cortical tpO_2_ decreases as a function of cortical depth (Feng et al., 1988). More recent work in rat has demonstrated deep cortical layers to possess longer and more heterogeneous microvascular transit times, suggestive of reduced oxygen extraction efficacy, in contrast to layer 4 transit times which were comparatively shorter and homogeneous, in keeping with optimal oxygen extraction (Jespersen and Østergaard, 2012; Merkle and Srinivasan, 2016). As a result, recurrent seizures with deep cortical epicenters may be particularly vulnerable to microregional tissue hypoxia given a comparatively low oxygen environment that is compounded by sub-optimal local oxygen extraction efficacy as well as a reduced microvasculature density (Masamoto et al., 2004). This recipe for hypoxia may therefore play a key role in shaping the deleterious effects of status epilepticus and epilepsia partialis continua, particularly in epilepsy syndromes associated with cortical laminar disruption, such as focal cortical dysplasia. Furthermore, it may play an important role in driving hypoxia-induced angiogenesis (Krock et al., 2011) and cerebrovascular remodeling that is frequently observed in epilepsy (Marchi and Lerner-Natoli, 2013).

In contrast, recurrent seizures with middle layer (L4-L5) epicenters were associated with increased cross laminar LFP synchrony, most likely as a consequence of travelling waves of ictal excitatory synaptic activity swelling into adjacent cortical laminae (Smith et al., 2016). Given the close correspondence between synaptic activity and hemodynamic signals (Logothetis et al., 2001), and recent reports of laminar dependent hemodynamic responses (Poplawsky et al., 2015), such seizures may therefore be able to maintain a tissue oxygen surplus by propagatively activating and availing themselves of local laminar neurovascular mechanisms, and exploiting a cortical angioarchitecture that is already optimized and accustomed to the elevated metabolic needs of the cortical input layers; such as increased microvascular (Masamoto et al., 2004) and arteriole primary branch density (Blinder et al., 2013), and red blood cell content and flow changes (Srinivasan and Radhakrishnan, 2014).

While it is as yet unclear why LFP synchrony between supragranular and underlying laminae is preferentially correlated to changes in tpO_2_ during the onset of recurrent seizures, it is interesting to note that GABAergic inhibitory neurons densely populate the superficial cortex (Meyer et al., 2011) and could play a key role in local neurovascular communication in the upper layers were it to be present. Our future work will address this directly through multi-depth electrophysiology and tpO_2_ recordings, and multi-laminar optogenetic activation of neuronal populations.

### 4.2 Preserved neural-hemodynamic and CBF-Hbt coupling

Our results have also shown that the relationship between neural activity and perfusion-based measures (i.e. CBF and Hbt) in whisker barrel cortex is linear, preserved, and remarkably robust during functional activation by sensory stimulation and intense recurrent seizures. This finding is significant as it provides support for the assumption of linearity in neural-hemodynamic coupling that is routinely made when interpreting perfusion-based neuroimaging signals, such as BOLD fMRI, in terms of underlying neural activity (Logothetis et al., 2001). This is additionally relevant given the rise of simultaneous clinical EEG-fMRI studies to localize hemodynamic correlates of electrophysiological epileptic activity during pre-surgical work-ups, and which are being increasingly conducted during ictal, rather than interictal periods, since this may provide a more accurate identification of the epileptogenic zone (Salek-Haddadi et al., 2002; Tyvaert et al., 2008). However, it is important to note that the observed linear regression slopes, while providing an excellent approximation, were somewhat steeper in the recurrent seizure condition compared to stimulation. This suggests that neural-CBF/Hbt coupling may be intrinsically non-linear at pathological extremes, a trait that may be more discernable during dynamic changes at seizure onset and offset (Ma et al., 2012; Harris et al., 2014a).

Similarly, we also found that that the relationship between CBF and Hbt (i.e. CBV) across all experimental conditions was well approximated by a linear function that was comparable along a continuum of focal to global hemodynamic changes. These similarities indicate that the flow-volume relationship is consistent irrespective of the mechanisms by which changes in flow are induced, and are of relevance to the calibration of the BOLD contrast in quantitative fMRI (Davis et al., 1998; Kida et al., 2007). This finding confirms and extends our previous work showing equivalent Grubb flow-volume coefficients during sensory stimulation and hypercapnia (Jones et al., 2001), and that of linearity between flow and volume during initiation and evolution of singular ictal-like discharges using non-concurrent techniques (Zhao et al., 2009). Notwithstanding, magnitude-related models do not capture temporal dynamics of flow and volume changes, with previous work confirming that Hbt/CBV onsets and returns to baseline more slowly than CBF (Figure 1 and 5; Jones et al., 2001; Kida et al., 2007), due to factors such as the viscoelastic nature of blood vessels (Mandeville et al., 1999) and/or capillary hyperemia (Hillman et al., 2007). Further studies using dynamic biophysical models are therefore needed to confirm that temporal flow-volume differences are not exacerbated under pathological conditions.

## 5. Conclusion

In summary, we have demonstrated, using a novel multi-modal approach, that laminar patterns of seizure initiation and evolution, evinced by translaminar synchrony, play an important role in shaping ictal cerebrovascular responses, and that these can be associated with prolonged periods of tissue hypoxia despite profound increases in perfusion and conserved neural-hemodynamic coupling. These findings provide new insights into the neurophysiogic correlates of seizure-related cerebrovascular responses, and reveal a hitherto unknown differential laminar susceptibility to tissue hypoxia during ictal events. It is thus tempting to speculate that laminar-specific predisposition to cerebrovascular dysfunction may play an important role in shaping the deleterious effects of recurrent seizures in epilepsy syndromes associated with laminar dysfunction, such as focal cortical dysplasia. However, future research, perhaps using simultaneous intracranial depth EEG and neuroimaging, is needed to confirm these findings in the clinical setting, and determine whether the observed mechanisms contribute to neurocognitive impairment in chronic epilepsy.

## Acknowledgements

We thank the technical staff of the University of Sheffield’s Department of Psychology, Dr Leonard Hetherington, Natalie Kennerley, Michael Port, and Llywelyn Lee.

## Funding

This work was supported by the Medical Research Council (grant number 141109); and Epilepsy Research United Kingdom (grant number 143100).

